# Colchicine Directly Targets Aldehyde Dehydrogenase 2 (ALDH2) to Suppress Radiation-Induced Senescence and Atherosclerosis

**DOI:** 10.64898/2026.05.25.726409

**Authors:** Jun-ichi Abe, Venkata SubrahmanyaKumar Samanthapudi, Weiqing Chen, Jonghae Lee, Ngoc Tuyet Tra, Gilbert F. Mejia, Oanh Hoang, Luis Antonio Rivera, Kun Yuan Chu, Masaki Osawa, Jung Hyun Kim, Shengyu Li, Kyung Ae Ko, Anilkumar K. Reddy, Silvia Fernanda López Moreno, Stefania Assunto Lenz, Keila Ostos Mendoza, Edgardo G. Sanchez, Anita Deswal, Joerg Herrmann, Keri L. Schadler, Laurent Yvan-Charvet, Charlotte Manisty, Pietro Ameri, Syed Wamique Yusuf, Rajneesh Pathania, Jared K. Burks, Nicolas L. Palaskas, Kevin T Nead, Michelle Hidebrandt, Clifton D. Fuller, Efstratios Koutrompakis, Sunil Krishnan, Steven H. Lin, Guangyu Wang, Nhat-Tu Le, Sivareddy Kotla

## Abstract

**Background:** Ionizing radiation (IR) accelerates atherosclerosis through induction of cellular senescence, DNA damage, defective efferocytosis, and dysregulation of clonal hematopoiesis (CH) drivers. Although low-dose colchicine reduces ischemic cardiovascular events in coronary artery disease, the precise molecular mechanisms underlying its vasculoprotective effects remain incompletely defined, and whether it mitigates radiation-associated vascular injury is unknown.

**Methods:** Bone marrow–derived macrophages (BMDMs) were pretreated with low-dose colchicine and exposed to 2 Gy IR. Molecular effects were assessed by RNA-seq, immunoblotting, and molecular docking. In vivo effects were tested in a partial carotid ligation (PLCL) model using spatial proteomics. Human monocyte-derived macrophages (HMDMs) from thoracic malignancy patients were analyzed before and after radiation therapy (RT).

**Results:** Low-dose colchicine suppressed IR-induced macrophage senescence signaling while preserving NRF2 activity. In a cell-free assay, colchicine directly activated aldehyde dehydrogenase 2 (ALDH2) in a dose-dependent manner (EC_50_ 1–5 nM), identifying ALDH2 as a direct molecular target of colchicine. Following irradiation, colchicine restored ALDH2, reduced mitochondrial (mt)ROS**-**dependent p90 ribosomal S6 kinase (p90RSK) activation and lipid peroxidation, preserved TET2 and DNMT3A expression, and rescued impaired efferocytosis while preventing nicotinamide adenine dinucleotide (NAD⁺) and adenosine triphosphate (ATP) depletion. These protective effects were ALDH2-dependent, as they were lost with ALDH2 inhibition or depletion and were mimicked by pharmacologic ALDH2 activation. In vivo, colchicine attenuated radiation-induced atherosclerosis and macrophage senescence-associated stemness (SAS). Consistently, macrophages from patients after RT showed reduced ALDH2 with increased mtROS, lipid peroxidation, and senescence.

**Conclusion:** These findings identify ALDH2 as a previously unrecognized molecular target of colchicine that links mitochondrial redox control to suppression of radiation-induced macrophage senescence and atherosclerosis and may contribute to the efficacy of low-dose colchicine in cardiovascular disease.

## Introduction

Radiation-induced cardiovascular disease (RICVD) is emerging as a critical issue among cancer survivors, warranting increased attention and research. Radiation therapy (RT) is a cornerstone in the management of primary thoracic malignancies, including lung cancer, breast cancer, and lymphoma. However, its delayed effects on the cardiovascular system contribute to significant long-term morbidity and mortality in this population^1^. Cardiovascular disease (CVD) is a leading cause of premature morbidity and mortality in survivors of breast cancer^2–4^, Hodgkin or non-Hodgkin lymphoma^5–8^, and lung cancer^9,10^, particularly more than five years after diagnosis and treatment. For example, patients with Hodgkin lymphoma who undergo RT to the chest face an approximately 7-fold increased risk of developing CVD, as demonstrated by multivariate analyses accounting for other risk factors^11^. Similarly, in breast cancer survivors, the incidence of major coronary events rises linearly by 7.4% for every Gray (Gy) of radiation exposure. Despite advances in modern RT techniques that reduce cardiac radiation exposure, breast cancer patients still often receive doses ranging from 1 to 5 Gy to the heart^1^. A deeper understanding of the mechanisms underlying RT-induced CVD is essential to developing targeted preventive and therapeutic strategies. Addressing this critical issue can significantly improve long-term outcomes for cancer survivors and mitigate one of the most pressing challenges in survivorship care.

CVD risk factors tend to be more pronounced in cancer survivors compared to the general population, with higher average levels of body mass index (BMI), C-reactive protein (CRP), and low-density lipoprotein (LDL) cholesterol. A notable and commonly observed phenotype in cancer survivors is accelerated cellular senescence^12–14^. Emerging evidence indicates that genotoxic chemotherapy agents and radiation contribute to this phenomenon by inducing mitochondrial dysfunction, inflammation, and cellular senescence, characterized by the senescence-associated secretory phenotype (SASP). DNA-damaging agents are known to trigger SASP within a relatively short timeframe, typically between 3 to 10 days, irrespective of significant telomere shortening^15^. While most DNA damage is efficiently repaired by the DNA damage response (DDR) mechanisms within 24 hours of the initial stress^16^, telomeric DNA damage exhibits a striking persistence, often lasting for months^17^. This prolonged telomeric DNA damage may provide a mechanistic basis for the late or delayed onset of CVD observed in cancer survivors following genotoxic therapies. The persistence of telomeric DNA damage and its associated induction of SASP underscore a plausible pathway linking cancer treatments to long-term cardiovascular complications. This connection highlights the importance of addressing and mitigating these late effects as part of survivorship care in oncology patients.

The benefits of colchicine in reducing atherosclerotic cardiovascular disease (ASCVD) events have been demonstrated through two pivotal clinical trials, the LoDoCo2 and COLCOT studies. Both trials evaluated the efficacy of adding low-dose colchicine (0.5 mg daily) to standard medical therapy, which included antiplatelet agents and statins, in mitigating major adverse cardiovascular events (MACE). The COLCOT trial focused on patients who had recently experienced a myocardial infarction (within 30 days prior to enrollment), whereas the LoDoCo2 trial targeted individuals with chronic coronary artery disease, defined as at least six months post-acute coronary syndrome. In both cohorts, the addition of low-dose colchicine to standard therapy significantly reduced the risk of MACE by more than 30% compared to placebo, highlighting its efficacy as a therapeutic adjunct in diverse cardiovascular contexts.

Mechanistically, colchicine exerts its effects by binding to tubulin, thereby disrupting microtubule-dependent processes in rapidly dividing cells. This disruption extends to the inhibition of inflammatory pathways, particularly through its impact on the NLRP3 inflammasome. Colchicine specifically reduces interleukin-1β secretion by preventing the colocalization of NLRP3 inflammasome components, namely ASC and NLRP3. Importantly, this targeted action does not affect other inflammasomes, such as NLRC4 or AIM2, suggesting that colchicine selectively modulates the inflammatory response. These findings underscore colchicine’s role in attenuating systemic inflammation triggered by molecular patterns that activate the NLRP3 inflammasome while preserving critical host defense mechanisms. This dual capability positions colchicine as a promising intervention for reducing ASCVD events by addressing underlying inflammatory pathways central to disease progression. However, the ability of colchicine to reduce SASP and impact RICVD remains unclear. While colchicine is known to target inflammation via the NLRP3 inflammasome, its effect on senescence—a key driver of RICVD—has not been fully studied.

Clonal hematopoiesis of indeterminate potential (CHIP) arises from somatic mutations in hematopoietic stem or progenitor cells, leading to clonal expansion without overt malignancy. Common mutations in CH driver genes such as *TET2*, *DNMT3A*, *ASXL1*, and *JAK2*—primarily loss-of-function—have been associated with a more than two-fold increased risk of cardiovascular disease (CVD), largely due to chronic inflammation mediated by mutant immune cells. Despite their clinical significance, the regulation of CH driver expression in the context of RICVD remains poorly understood. CHIP mutations are frequently observed in cancer survivors, particularly those exposed to radiotherapy, and may exacerbate vascular toxicity and impair tissue repair, contributing to late-onset complications such as RICVD. In this study, we demonstrate that colchicine not only reverses IR-induced suppression of CH drivers but also directly enhances the activity of ALDH2—a mitochondrial detoxifying enzyme whose activation by colchicine has not been previously reported. This novel finding reveals that ALDH2 plays a central role in mediating colchicine’s protective effects, as its inhibition abolishes the restoration of CH driver expression and the reduction of oxidative damage. These results position ALDH2 as a key regulatory node linking mitochondrial detoxification to clonal hematopoiesis and vascular inflammation, and establish it as a critical target in mitigating RICVD.

## METHODS

Details of the antibodies and reagents, experimental models and subject details, experimental procedures, imaging with COMET™ multi-fluorescence imaging and imaging mass cytometry (IMC), and power calculation are included in the online supplemental information.

### Data Sharing Availability

The data, analytic methods, and study materials that support the findings of this study are available in the Data supplement or from the corresponding authors upon reasonable request.

### Institutional Approvals

The study was approved by the Institutional Review Board of the University of Texas MD Anderson Cancer Center (#PA16-0971). All procedures adhered to applicable federal regulations, including the Health Insurance Portability and Accountability Act (HIPAA) and the Common Rule (45 CFR 46).

All animal studies were conducted in accordance with institutional and federal ethical guidelines and were approved by the Institutional Animal Care and Use Committees (IACUC) at The University of Texas MD Anderson Cancer Center (protocols 00001652 and 00001952).

### In vivo mouse model

We created an in-vivo mouse model to determine whether colchicine can inhibit IR-induced atherosclerosis lesions. We utilized our reported partial left carotid ligation (PLCL) model. This model was used to detect whether colchicine can inhibit IR-mediated acceleration of the atherosclerosis process. As shown in Fig. 7A, we applied AAV-PCSK9 and started a high-fat diet (HFD) for 13 days before initiating colchicine treatment (0.01 mg/kg/day, intraperitoneally). To control body weight loss in IR-exposed mice, we switched to a normal chow diet for two weeks after completing IR, then reverted to HFD. One week after resuming HFD, we performed PLCL and waited two weeks before isolating the carotid arteries. We found that cholesterol levels in both groups were significantly upregulated, with no difference between the colchicine and vehicle groups.

We evaluated the atherosclerosis lesions through gross evaluation and calculated the intima/media ratio after histological evaluation, as previously described.

### Binding Affinity and Interaction Analysis Following Molecular Docking

We investigated potential binding of colchicine to ALDH2 using a structure-based docking approach. The crystallographic coordinates of human ALDH2 were obtained from the Protein Data Bank: 3INJ (ALDH2–Alda-1 complex) and 4FR8 (apoenzyme). Each structure was prepared for docking using AutoDockTools (v1.5.7). All crystallographic waters, ions, and non-essential ligands were removed; polar hydrogens were added and Gasteiger charges assigned.

The receptors were kept rigid during all simulations. Ligand structures for colchicine (PubChem CID 6167) and CVT-10216 (PubChem CID 24797264) were downloaded from PubChem, protonated at physiological pH, and energy-minimized with the MMFF94 force field in Avogadro (v1.2.0). The resulting structures were saved in PDBQT format with all rotatable bonds defined.

Molecular docking was performed using AutoDock Vina v1.2.5 employing the default *vina* scoring function (Eberhardt *et al.*, 2021; Trott & Olson, 2010). For 3INJ, the docking grid was centered on the Alda-1 activator pocket (X = 6.365, Y = −27.38, Z = 44.879); for 4FR8, it was centered at X = 1.43, Y = 49.958, Z = 49.385. In both cases the grid size was 24 × 24 × 24 Å with 0.375 Å spacing and an exhaustiveness value of 128. Twenty poses were generated for each ligand. Docking was executed on a 64-bit Ubuntu 22.04 workstation equipped with an 8-core 3.8 GHz CPU and 16 GB RAM. Binding energies (kcal mol⁻¹) and RMSD values were taken directly from the Vina log files to assess convergence and pose clustering.

The best-scoring pose for each ligand–receptor pair was analyzed using PyMOL (v2.5) and Discovery Studio Visualizer (v21.1). Hydrogen-bond distances, hydrophobic contacts, and π–π stacking interactions were identified, and residues within 4 Å of each ligand were recorded.

Colchicine and CVT-10216 docking results were compared with the Alda-1–bound conformation in 3INJ to validate targeting of the activator (vestibule) pocket. Shared and unique contact residues were mapped to evaluate potential competitive binding within this site. Figures were rendered in PyMOL, and all distances are reported in Å.

### Statistical Analysis

All statistical analyses, including those applied to the IMC/COMET and spatial metabolomics datasets, are described in detail in the Supplemental Methods section.

All statistical analyses were performed using GraphPad Prism software version 9.0.0 or 10 (GraphPad Software, San Diego, CA, USA). The Shapiro–Wilk test was used to assess the normality of each dataset. For comparisons among more than two groups, ordinary one-way or two-way analysis of variance (ANOVA) was conducted, followed by Tukey’s post hoc test, provided the data met normality assumptions. For two-group comparisons with normally distributed data, unpaired two-tailed Student’s *t*-tests were applied. When data failed the Shapiro–Wilk normality test, the nonparametric Mann–Whitney *U* test was used for two-group comparisons, and the Kruskal–Wallis test followed by Dunn’s post hoc test was applied for multi-group comparisons. A *P* value < 0.05 was considered statistically significant, except for analyses involving COMET data. All experiments were performed independently unless otherwise indicated.

All data visualizations, including violin plots and bar graphs, were generated using GraphPad Prism version 10 (GraphPad Software, San Diego, CA, USA). Detailed statistical information—including sample sizes (*n*), normality test outcomes, statistical methods used, and *P* values for each figure—is provided in Table S1.

### COMET Data Analysis

Image files from COMET (a sequential immunofluorescence platform) were exported as OME-TIFF and MCD (mass cytometry data) formats, respectively. These files were imported into Visiopharm software and aligned using the TissueAlign module with a U-net-based approach to generate a unified image database for high-dimensional single-cell analysis.

All data visualizations, including violin plots and bar graphs, were generated using GraphPad Prism version 10 (GraphPad Software, San Diego, CA, USA). Detailed statistical information—including sample sizes (*n*), normality test outcomes, statistical methods used, and *P* values for each figure—is provided in Table S1.

**Supplementary Table 1A (Main figures), 1B (Supplementary figures), and 1C (Spatial data):** Statistical analysis for each experiment.

**Supplementary Table 2:** Controls for each relative value.

**Supplementary Table 3:** Antibodies for COMET

**Supplementary File 1:** Spatial proteomics intensity datasets used for analyses in Fig. 7, Fig. 8A, and Supplementary Figs. S2–S4 from the mouse plaque study, including immune cells, endothelial cells (ECs), and smooth muscle cells (SMCs).

**Supplementary File 2:** Quantification of distances between CD31⁺ endothelial cells and F4/80⁺ macrophages used in Fig. 7L.

**Supplementary File 3:** Quantification of distances between CD4⁺, CD8⁺, and CD45⁺ cells and F4/80⁺ macrophages used in Fig. 7K.

## Results

### Colchicine Restores NRF2 Activity and Reverses IR-Induced Transcriptomic Changes in Macrophages

To investigate the effects of IR on macrophage function, bone marrow-derived macrophages (BMDMs) were isolated and exposed to 2 Gy of IR. This dose was selected based on prior dose-response experiments, which showed that 2 Gy significantly reduced succinate dehydrogenase activity without inducing apoptosis—allowing us to examine persistent cellular effects rather than acute cell death^18^. BMDMs were pretreated with colchicine or vehicle for 1 hour prior to IR exposure, and RNA sequencing was performed 24 hours post-irradiation. Gene ontology (GO) enrichment analysis revealed that IR significantly altered biological processes associated with the apoptotic signaling pathway (positive regulation of apoptotic processes), the cell migration pathway (positive regulation of cell migration), and the cell survival pathway (negative regulation of apoptotic processes) (**Fig. 1A, C, E**). Colchicine pretreatment modulated these IR-induced transcriptomic changes, with the top enriched biological processes corresponding to the inflammatory response pathway (regulation of inflammatory processes), the cell migration pathway (positive regulation of cell migration), and the apoptotic signaling pathway (positive regulation of apoptotic processes) (**Fig. 1B, D, and F**).

**Figure 1.**
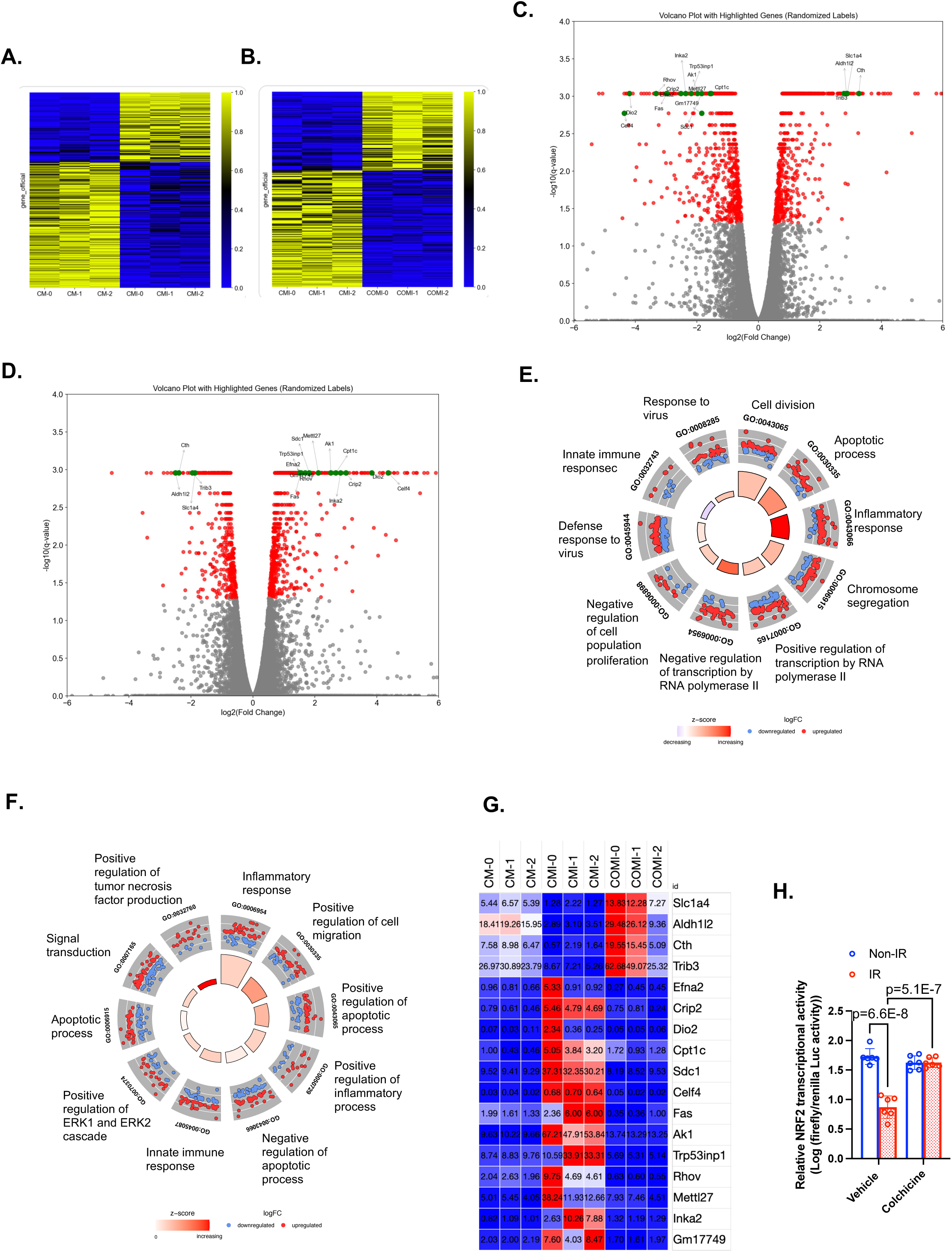
Colchicine modulates ionizing radiation–induced transcriptional reprogramming and restores NRF2 signaling in macrophages. Bone marrow–derived macrophages (BMDMs) were pretreated with colchicine or vehicle for 1 hour and exposed to 2 Gy ionizing radiation (IR). RNA sequencing was performed 24 hours after IR exposure to assess transcriptomic changes. **(A–B)** Heatmaps showing differentially expressed genes in BMDMs following IR exposure (**A**) and colchicine pretreatment with IR (**B**), illustrating distinct transcriptional reprogramming patterns. **(C–D)** Volcano plots of RNA-seq data highlighting significantly upregulated and downregulated genes in IR-treated macrophages (**C**) and colchicine-pretreated macrophages following IR (**D**). **(E–F)** Gene ontology (GO) enrichment analyses. IR significantly modulated pathways related to apoptotic signaling, cell migration, and survival (**E**), whereas colchicine altered IR-responsive pathways, including regulation of inflammatory response, cell migration, and apoptotic signaling (**F**). **(G)** Heatmap of 17 genes whose IR-induced expression changes were significantly reversed by colchicine, including NRF2- and ATF4-regulated genes such as *Slc1a4*, *Aldh1/2*, *Cth*, and *Trib3*. **(H)** Quantification of NRF2 transcriptional activity demonstrating suppression by IR and restoration with colchicine pretreatment.

To further understand colchicine’s role, we identified 17 genes whose IR-induced expression changes were significantly reversed by colchicine (**Fig. 1G**). Among these, *Slc1a4*^19,20^, *Aldh1/2*^21^, *Cth*^22^, and *Trib3*^23^ are known targets of the transcription factors ATF4 and NRF2. While ATF4 is typically upregulated by IR^24^, our previous findings indicate that NRF2 is downregulated^18^. Focusing on NRF2, we found that IR significantly suppressed its transcriptional activity, whereas colchicine pretreatment restored it (**Fig. 1H**). These results suggest that colchicine may counteract IR-induced suppression of the NRF2-mediated oxidative stress response pathway, thereby regulating the expression of key downstream genes involved in cellular stress adaptation and survival.

### Colchicine Attenuates Ionizing Radiation–Induced Cellular Senescence in Macrophages

Previous studies suggest that colchicine can inhibit SIRT2, a NAD⁺-dependent deacetylase, and suppress inflammasome activation^25^. Given the established role of NAD⁺ depletion and SIRT2 in regulating the senescence-associated secretory phenotype (SASP), we investigated whether colchicine modulates ionizing radiation (IR)–induced inflammation and senescence in macrophages. We first examined activation of the NF-κB signaling pathway and inflammatory cytokine secretion. IR exposure induced robust NF-κB activation, and colchicine treatment significantly attenuated NF-κB activation, although this inhibition was partial (**Fig. 2A**). Consistent with this interpretation, combined colchicine and IR treatment was associated with upregulation of inflammatory response genes at the transcriptomic level (**Fig. 1F**).

**Figure 2.**
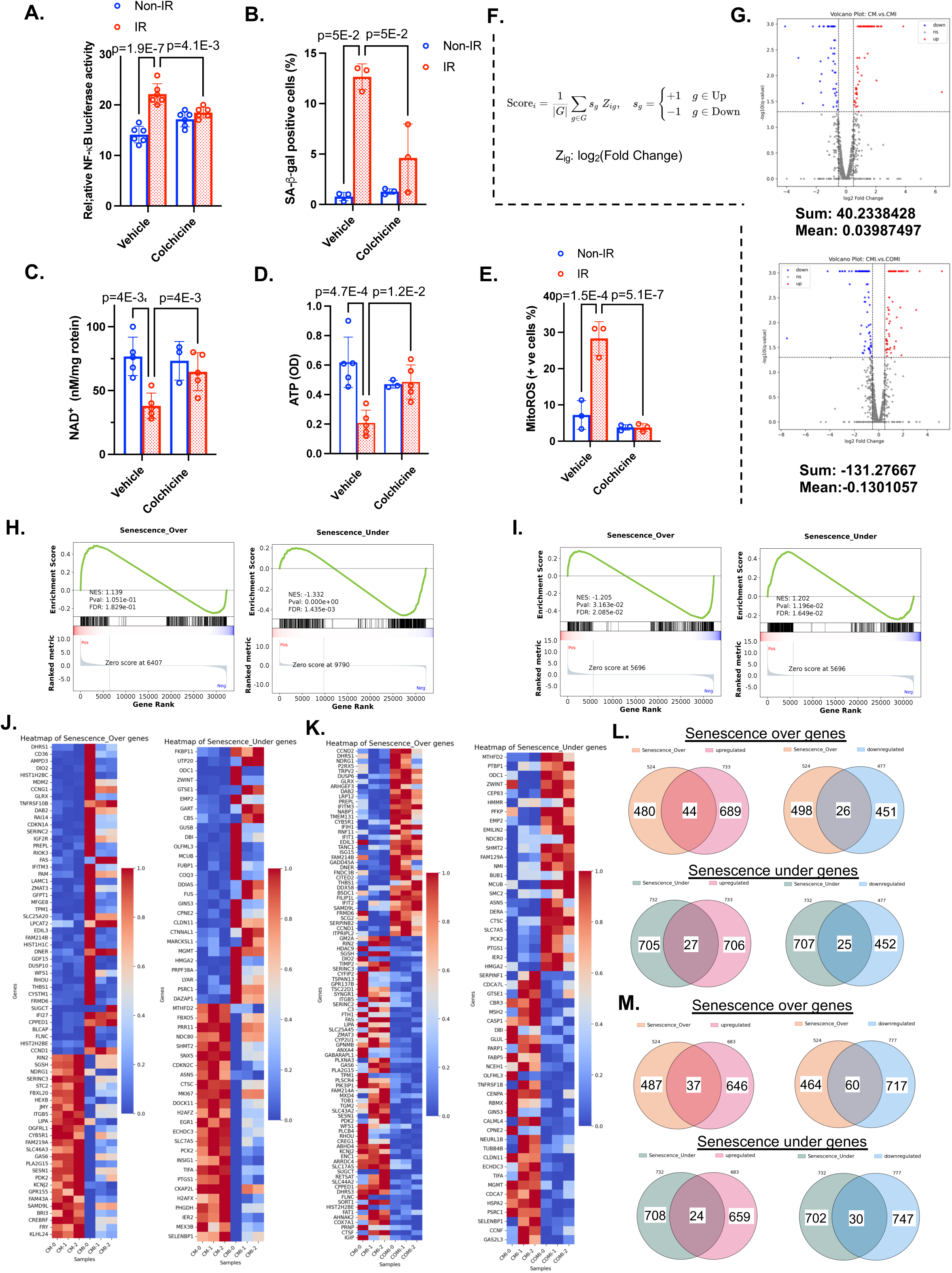
Colchicine suppresses IR-induced senescence signaling in macrophages. BMDMs were pretreated with colchicine or vehicle and exposed to ionizing radiation (IR, 2 Gy). **(A)** NF-κB transcriptional activity was measured by luciferase reporter assay 12 hours post-irradiation. **(B–E)** Cellular senescence and metabolic alterations. **(B)** Senescence-associated β-galactosidase (SA-βgal) staining was used to quantify senescent cells. **(C, D)** Intracellular NAD⁺ and ATP levels were measured 24 hours after IR. **(E)** Mitochondrial reactive oxygen species (mtROS) levels were assessed using MitoSOX staining. **(F, G)** Senescence-associated genes from the CellAge database (n = 1,009) were used to assess radiation-induced transcriptional changes. A composite senescence score was calculated by averaging gene-level Z-scores, with positive values indicating upregulation and negative values indicating downregulation. Ionizing radiation increased the composite senescence score, reflecting global activation of a senescence-associated transcriptional program, as shown by differential gene expression in the volcano plot comparing vehicle control and irradiated samples (CM vs CMI, top). In contrast, colchicine treatment shifted the composite score to a negative value, indicating overall suppression of senescence-associated gene expression, as illustrated by the volcano plot comparing irradiated samples with or without colchicine (CMI vs COMI, bottom). **(H-I)** Gene set enrichment analysis (GSEA) of senescence-associated gene sets. Panel I shows that ionizing radiation activates a senescence-associated transcriptional program, with enrichment of genes typically upregulated during senescence (Senescence_Over; NES = 1.139) and significant suppression of genes normally repressed during senescence (Senescence_Under; NES = –1.332, FDR = 0.0014). Panel J demonstrates that colchicine reverses these radiation-induced changes, with negative enrichment of Senescence_Over genes (NES = –1.205, FDR = 0.021) and positive enrichment of Senescence_Under genes (NES = 1.202, FDR = 0.016), consistent with attenuation of radiation-induced senescence. **(J-K)** Heatmaps of senescence-associated genes, with genes typically upregulated during senescence (Senescence_Over, left) and downregulated during senescence (Senescence_Under, right). Panel K shows radiation-induced senescence-associated transcriptional changes, whereas panel L shows reversal of these changes with colchicine treatment. **(L-M)** Venn diagrams of differentially expressed senescence-associated genes, with Senescence_Over genes shown at the top and Senescence_Under genes shown at the bottom. Panel M depicts senescence-related genes altered by ionizing radiation, while panel N shows genes modulated by colchicine in irradiated samples. Numbers indicate shared and condition-specific genes.

We next assessed cellular senescence. IR significantly increased senescence-associated β-galactosidase (SA-β-gal) activity and depleted intracellular NAD⁺ levels, effects that were markedly reversed by colchicine pretreatment (**Fig. 2B, C**). Because ATP production is tightly coupled to NAD⁺ availability, we also measured cellular ATP levels and found that colchicine significantly restored ATP levels that were otherwise reduced following IR exposure (**Fig. 2D**). Together, these findings indicate that colchicine protects against IR-induced inflammatory activation and metabolic dysfunction associated with cellular senescence. Lastly, given the proposed role of mitochondrial reactive oxygen species (mtROS) in NAD⁺ depletion and SASP induction^18^, we measured mtROS production following IR and found that colchicine effectively suppressed IR-induced mtROS generation (**Fig. 2E**). Together, these results suggest that colchicine mitigates IR-induced senescence and inflammation in macrophages by preserving NAD⁺/ATP levels and inhibiting mtROS production and NF-κB activation.

To further investigate the effect of colchicine on IR–induced senescence in macrophages, we analyzed transcriptomic signatures of senescence induction. Radiation exposure significantly altered the expression of senescence-associated genes curated from the CellAge database. A composite senescence score was calculated for each sample by averaging the Z-scores of 1,009 senescence-related genes, with gene directionality defined by their response to radiation (+1 for radiation-upregulated genes and −1 for radiation-downregulated genes) (**Fig. 2F**). The summed senescence score was 40.2, with a mean score of 0.03987, indicating an overall increase in radiation-induced senescence-associated transcriptional activity (**Fig. 2F, G, top**).

Upon gene set enrichment analysis (GSEA), radiation exposure was found to induce a transcriptional signature consistent with cellular senescence. Specifically, genes typically upregulated during senescence (Senescence_Over) exhibited a positive enrichment trend in irradiated samples compared to non-irradiated controls (NES = 1.139, FDR = 0.183), indicating partial activation of the senescence-associated program. In contrast, genes normally repressed during senescence (Senescence_Under) showed a significant negative enrichment following radiation (NES = –1.332, FDR = 0.0014), reflecting strong downregulation of anti-senescence and cell cycle–related pathways. Together, these findings suggest that ionizing radiation promotes a robust senescent phenotype characterized by coordinated activation of SASP-related genes and suppression of proliferation-associated transcriptional networks (**Fig. 2H**).

Gene Ontology (GO) pathway enrichment analysis revealed marked transcriptional reprogramming following ionizing radiation. The GO-derived heatmap displayed senescence-over and senescence-under gene sets separately, showing coordinated activation of senescence-over genes and repression of senescence-under genes in irradiated samples (**Fig. 2J**). **Table 1** (upper) quantitatively confirmed statistically significant enrichment changes in both senescence-over and senescence-under gene sets. Venn diagram analysis further illustrated these effects by independently showing overlap between radiation-regulated genes and senescence-over or senescence-under gene sets, respectively (**Fig. 2L**). Together, these analyses indicate that radiation induces a robust and coordinated senescence-associated transcriptional response.

Comparison of vehicle plus radiation (CMI) and colchicine plus radiation (COMI) revealed that colchicine broadly suppresses radiation-induced cellular senescence. Analysis of the CellAge senescence-associated gene set showed a negative composite senescence score in the COMI group, indicating an overall reduction in senescence-related transcriptional activity compared with CMI (**Fig. 2G, bottom**). Consistently, GSEA demonstrated reduced expression of senescence-over genes and restoration of senescence-under genes in COMI relative to CMI, indicating reversal of radiation-induced senescence-associated transcriptional programs (**Fig. 2I**).

GO pathway enrichment analysis further confirmed that colchicine counteracts radiation-driven transcriptional reprogramming, suppressing senescence- and inflammation-related pathways while promoting pathways associated with metabolic recovery and cellular homeostasis (**Fig. 2K**, **Table 1 (lower), Fig.2M**). Together, these findings demonstrate that colchicine effectively inhibits IR-mediated senescence across gene-, gene set–, and pathway-level analyses.

### Colchicine Prevents IR-Induced ALDH2–4-HNE Dysregulation and Loss of CH Driver Genes Through Modulation of the mtROS–p90RSK Feedback Loop

Although DNA methylation has been implicated in the regulation of SASP^26^, the specific contribution of clonal hematopoiesis (CH) driver genes such as TET2 and DNMT3a to IR-induced SASP has not been well defined. To address this, we examined the expression of CH drivers after IR exposure and assessed the modulatory effects of colchicine. Our results showed that IR significantly reduced TET2 and DNMT3a protein expression, whereas pretreatment with colchicine effectively prevented this downregulation (**Fig. 3A, B**). Interestingly, IR did not alter TET2 or DNMT3a mRNA expression (**Fig. 3C**), suggesting that these protein changes are likely regulated at the post-transcriptional level rather than through transcriptional control.

**Figure 3.**
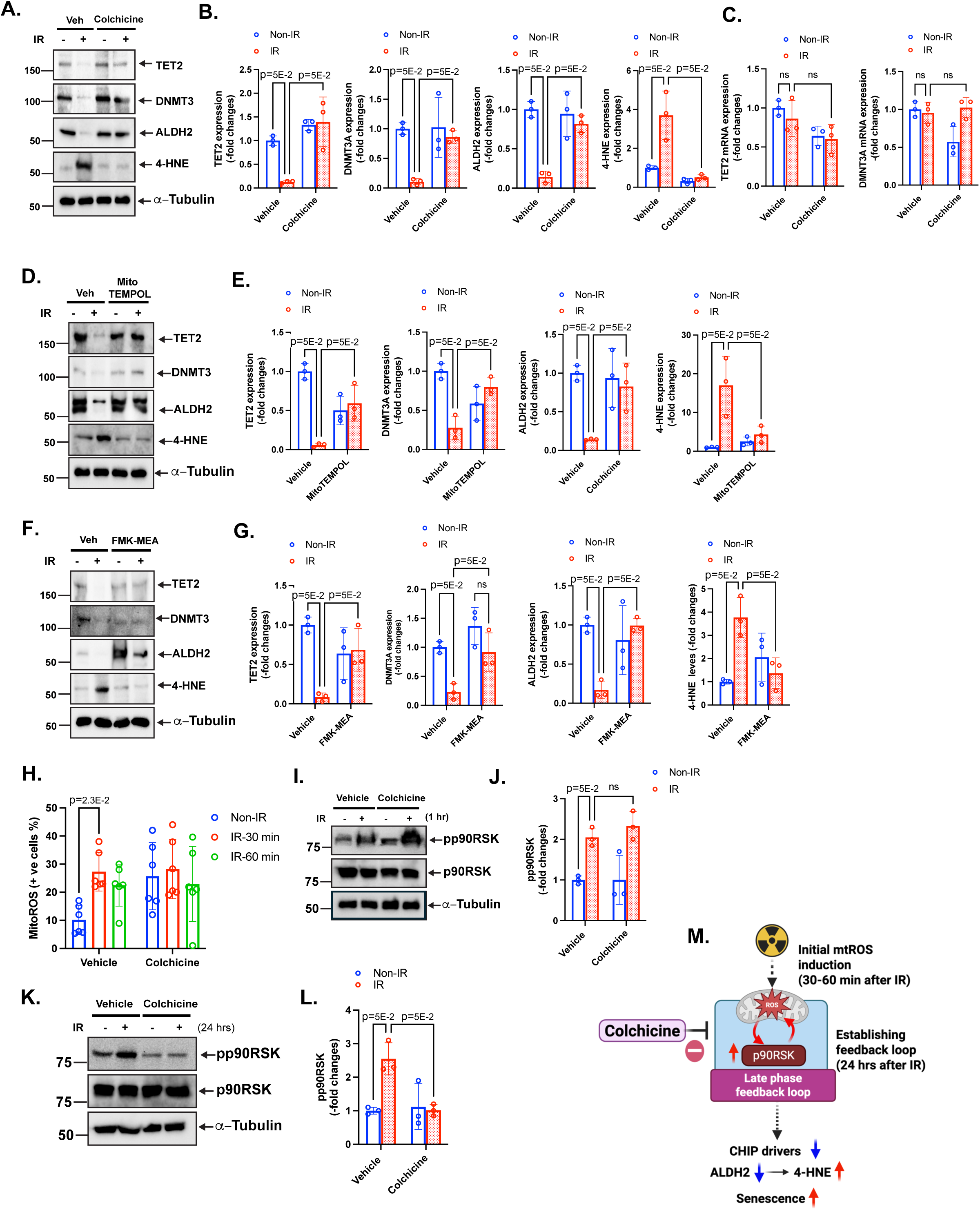
Colchicine disrupts the IR-induced mtROS–p90RSK feedback loop and preserves CH driver expression via ALDH2 signaling. **(A, B, D-G)** Effects of pharmacological modulation of mitochondrial ROS and downstream signaling. BMDMs were pretreated with colchicine (**A, B**), the mitochondrial ROS scavenger MitoTEMPO (**D–E**), or the p90RSK inhibitor FMK-MEA (**F–G**) prior to IR exposure. Protein expression was analyzed by Western blotting 24 hours post-irradiation (**B, D, F**), with corresponding densitometric quantification from three independent experiments (**C, E, G**). **(C)** mRNA expression of clonal hematopoiesis (CH) drivers (*TET2* and *DNMT3A*) in BMDMs 24 hours after ionizing radiation (IR), assessed by RT-qPCR. **(H)** Mitochondrial ROS (mtROS) levels measured by MitoSOX staining in BMDMs pretreated with colchicine and exposed to IR, demonstrating attenuation of IR-induced oxidative stress. **(I–L)** Time-dependent regulation of p90RSK activation. BMDMs were pretreated with colchicine or vehicle prior to IR exposure. p90RSK activity was assessed by Western blotting at early (1 hour; **I–J**) and late (24 hours; **K–L**) time points, with corresponding quantification. **(M)** Schematic illustrating that IR induces an early mtROS burst that activates p90RSK and establishes a late-phase mtROS–p90RSK positive feedback loop, resulting in suppression of CH drivers, ALDH2 downregulation, increased 4-HNE, and cellular senescence. Colchicine disrupts this loop, thereby preserving ALDH2 function, limiting lipid peroxidation, and attenuating senescence.

To further understand colchicine’s role, we identified 17 genes whose IR-induced expression changes were significantly reversed by colchicine (**Fig. 1G**), including Slc1a4, Aldh1/2, Cth, and Trib3, known ATF4/NRF2 targets. Given the central role of ALDH2 in clearing 4-hydroxynonenal (4-HNE), limiting mitochondrial oxidative stress, and restraining vascular cellular senescence^27^, we next investigated whether modulation of this pathway underlies the protective effects of colchicine. Because impaired 4-HNE clearance promotes mtROS generation and p90RSK activation, as we previously reported^18^, restoration of Aldh1/2 expression suggested a potential mechanism for colchicine’s protective effects. Consistent with this, IR significantly reduced ALDH2 expression and increased 4-HNE accumulation, supporting activation of the ALDH2–4-HNE–mtROS–p90RSK axis (**Fig. 3A,B**).

Notably, pretreatment with colchicine, mitoTEMPOL (an mtROS-specific inhibitor), or FMK-MEA (a selective p90RSK inhibitor) reversed these effects, restoring ALDH2 expression and suppressing 4-HNE accumulation (**Fig. 3D-G**). We previously demonstrated that mtROS–p90RSK signaling forms a positive feedback loop following IR in macrophages, leading to macrophage senescence^18^. Together, these findings suggest that IR-induced ALDH2 downregulation, subsequent 4-HNE accumulation, and depletion of TET2 and DNMT3a are regulated through an mtROS–p90RSK–mediated positive feedback loop (**Fig. 3M**). Therefore, we hypothesized that colchicine inhibits mtROS-dependent p90RSK activation at an early stage following IR, thereby preventing ALDH2–4-HNE dysregulation and the reduction of TET2 and DNMT3a. However, colchicine did not affect IR-induced mtROS production or p90RSK activation within 1 hour after irradiation (**Fig. 3H-J**). In contrast, p90RSK activation at 24 hours post-IR was significantly attenuated by colchicine pretreatment (**Fig. 3K, L**). These findings suggest that initial mtROS-mediated p90RSK activation (30-60 min after IR) is unlikely to be a direct target of colchicine (**Fig. 3M**).

### ALDH2 Activation Is Required for Colchicine-Mediated Suppression of the Senescence

Because RNA-seq indicated recovery of ALDH1/2 expression by colchicine following IR-induced suppression (**Fig. 1G**), we next validated this effect by qPCR. Consistent with these findings, colchicine restored ALDH2, but not ALDH1, mRNA expression 24 hours after irradiation. Given that mtROS–p90RSK signaling may not represent the primary molecular target of colchicine, we next investigated whether ALDH2 activation contributes to colchicine-mediated suppression of the senescence after IR. To address this, we first examined the effects of Alda-1, a selective ALDH2 activator. Pretreatment with Alda-1 markedly decreased SA-β-gal activity, and prevented NAD⁺ depletion and mtROS accumulation (**Fig. 4B**). Alda-1 attenuated IR-induced ALDH2 reduction and 4-HNE accumulation, which was associated with preservation of TET2 and DNMT3A expression and reduced activation of p90RSK (**Fig. 4C,D**). In parallel, ALDA-1 blunted induction of p53 and p21, prevented Lamin B1 loss, and maintained Trx expression (**Fig. 4E,F**), consistent with mitigation of aldehyde-driven redox stress and attenuation of p53-associated senescence.

**Figure 4.**
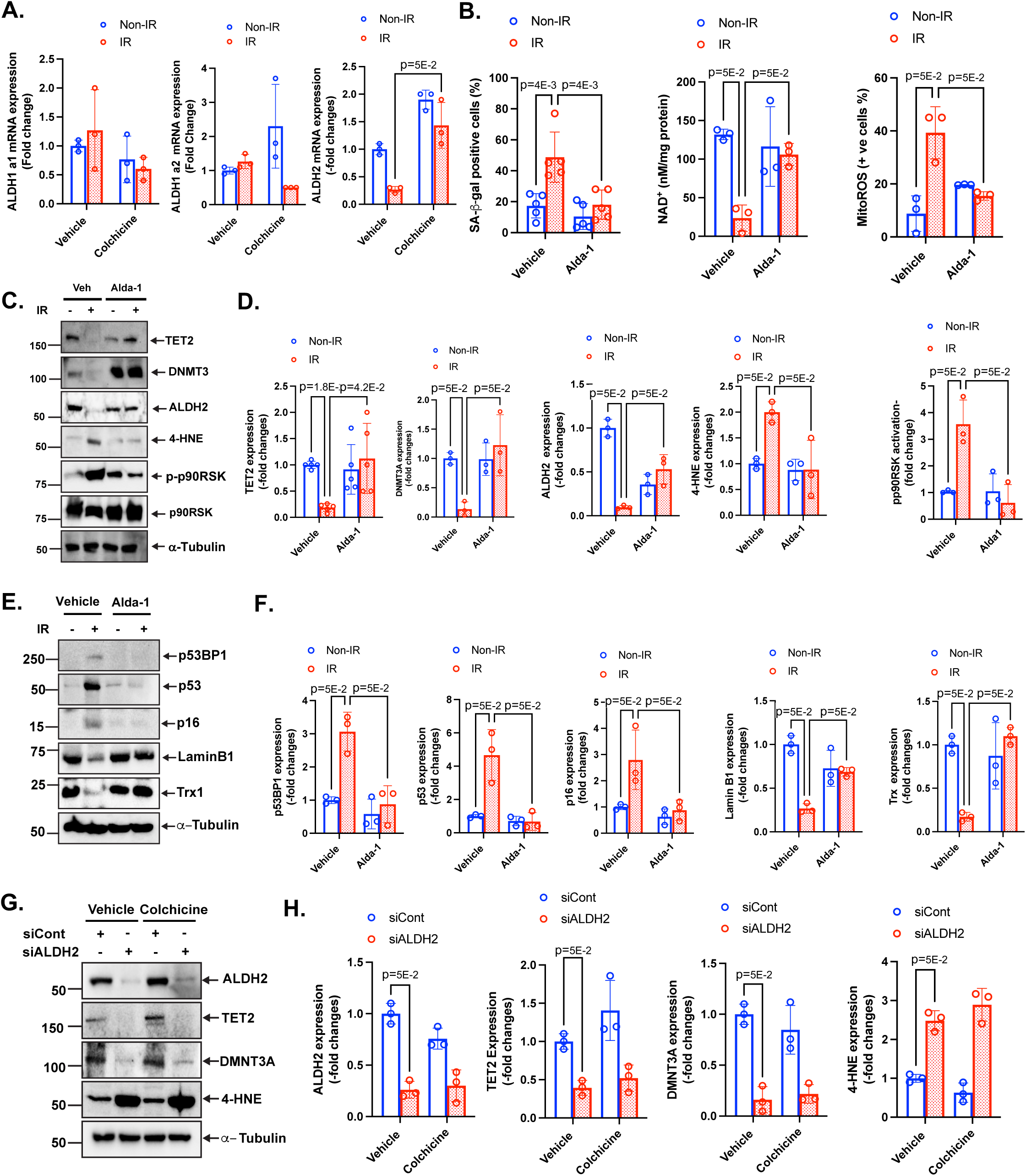
ALDH2 specific activator (Alda-1) attenuates IR-induced mtROS-p90RSK positive feedback loop, reduction of CH drivers, ALDH2-4HNE signaling, and senescence. **(A)** BMDMs were pretreated with colchicine or vehicle, followed by exposure to IR. ALDH1 and 2 mRNA levels were assessed after 24 hrs by using ALDH1 and 2 specific primers. **(B-F)** BMDMs were pretreated with Alda-1 or vehicle, followed by exposure to IR (2 Gy). SA-βgal, NAD^+^, and mtROS level were measured (**B**), and protein expression was assessed by Western blotting after 24hrs using the indicated antibodies (**C, E**), and quantification (**D, F**). **(G, H)** BMDMs were pretreated with ALDH2 or control siRNA, followed by colchicine or vehicle. After 1 h, cells were exposed to IR. Protein expression was assessed by Western blotting after 24hrs using the indicated antibodies (**G**), and quantification (**H**).

To determine whether the effects of colchicine are dependent on ALDH2, ALDH2 expression was depleted by siRNA prior to colchicine treatment (**Fig. 4G**). ALDH2 knockdown alone resulted in a significant reduction in TET2 and DNMT3A protein levels and a concomitant increase in 4-HNE accumulation, indicating enhanced aldehyde stress. Under these conditions, colchicine failed to restore TET2 or DNMT3A expression or to reduce 4-HNE levels. These findings indicate that ALDH2 is required for colchicine-mediated preservation of epigenetic regulators and suppression of aldehyde stress (**Fig. 4G, H**).

Next, to determine whether colchicine acts through ALDH2, BMDMs were pretreated with the ALDH2 inhibitor CVT-10216 before colchicine exposure and subsequent IR (**Fig. 5A**). As expected, colchicine reversed IR-induced reductions in TET2, DNMT3A, and ALDH2 expression and reduced 4-HNE accumulation. However, CVT-10216 pretreatment abolished these effects, as colchicine no longer attenuated IR-induced 4-HNE accumulation or restored TET2 and DNMT3A expression (**Fig. 5A, B**). Consistently, the ability of colchicine to prevent IR-induced NAD⁺ depletion was significantly inhibited by CVT-10216 (**Fig. 5C**). In parallel, ALDH2 knockdown using siRNA abolished colchicine-mediated attenuation of ALDH2 depletion–associated senescence markers, including accumulation of 53BP1 and p16 and reduction of Lamin B1 (**Fig. 5D**). Furthermore, colchicine failed to suppress SA-β-gal induction in endothelial cells following ALDH2 knockdown (**Fig. 5E**). Together, these findings indicate that ALDH2 activity is required for colchicine-mediated protection against IR-induced aldehyde stress, epigenetic dysregulation, and cellular senescence.

**Figure 5.**
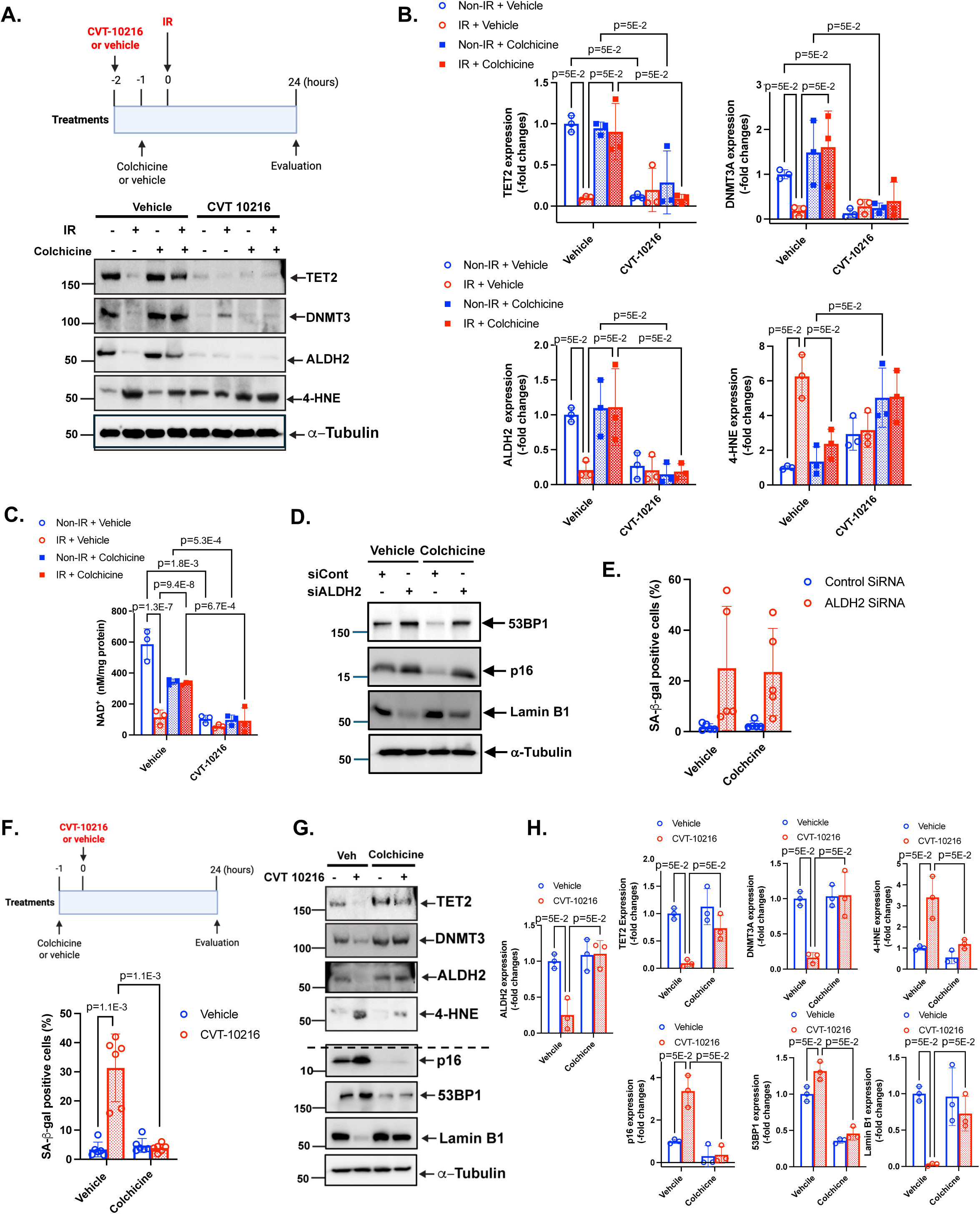
ALDH2 specific inhibitor (CVT-10216) attenuates IR-induced mtROS-p90RSK positive feedback loop, reduction of CH drivers, ALDH2-4HNE signaling, and senescence. **(A-C)** BMDMs were pretreated with CVT-10216 or vehicle, followed by colchicine or vehicle. After 1 h, cells were exposed to IR. Protein expression was assessed by Western blotting 24 h after IR using the indicated antibodies (**A**), with densitometric quantification (**B**). Intracellular NAD⁺ levels were measured (**C**). **(D, E)** BMDMs were pretreated with ALDH2 or control siRNA, followed by colchicine or vehicle. After 1 h, cells were exposed to IR. Protein expression was assessed by Western blotting after 24hrs using the indicated antibodies (**D**), and quantification (**Fig. S1**), and SA-βgal (**E**) was measured. (**F-H**) BMDMs were pretreated with cholchicine or vehicle, followed by CVT-10216 or vehicle treatment. SA-βgal (**F**) and protein expression was assessed by Western blotting after 24hrs using the indicated antibodies (**G**), and quantification (**H**).

Next, to determine whether colchicine modulates cellular senescence through preservation of ALDH2 activity, BMDMs were pretreated with colchicine or vehicle prior to exposure to the ALDH2-specific inhibitor CVT-10216 (**Fig. 5F**). Treatment with CVT-10216 alone resulted in marked ALDH2 depletion, increased 4-HNE accumulation, reduced expression of TET2 and DNMT3A, and induction of senescence markers, including increased p16 and 53BP1 expression and loss of Lamin B1 (**Fig. 5G, H**). Notably, colchicine pretreatment significantly attenuated CVT-10216–induced SA-β-gal activity (**Fig. 5F**). Consistently, colchicine completely reversed CVT-10216–induced ALDH2 loss, suppressed 4-HNE accumulation, restored TET2 and DNMT3A expression, and mitigated senescence marker induction (**Fig. 5G, H**). Collectively, these findings indicate that colchicine counteracts ALDH2 inhibition–induced aldehyde stress, epigenetic dysregulation, and cellular senescence, supporting a critical requirement for intact ALDH2 activity in mediating the anti-senescent effects of colchicine. Moreover, the distinct outcomes observed under pretreatment (**Fig. 5F**) versus post-inhibition (**Fig. 5A**) conditions suggest that colchicine may directly modulate ALDH2 function rather than acting solely downstream of aldehyde stress.

### Differential Engagement of Colchicine and CVT-10216 with the Alda-1 Activator Pocket of ALDH2

To assess whether colchicine directly interacts with ALDH2, we performed *in silico* molecular docking analyses. Docking of colchicine to ALDH2 suggested a preferential association with the activator (vestibule) pocket, corresponding to the Alda-1 binding site observed in the 3INJ crystal structure. As shown in **Fig. 6A**, colchicine fits deeply within a hydrophobic cleft formed by Lys127, Asp147, Phe292, Asp457, and Phe459, anchoring through a hydrogen bond with Lys127 (2.4 Å) and stabilized by π–π interactions with Phe292 and Phe459. This region spatially overlaps with the Alda-1 binding cavity, which lies at the subunit interface adjacent to the substrate entrance tunnel. The similarity in binding location suggests that colchicine may interact with the same allosteric pocket utilized by Alda-1, potentially influencing the conformational dynamics of ALDH2 through partial occupation of the activator site rather than the catalytic core.

**Figure 6.**
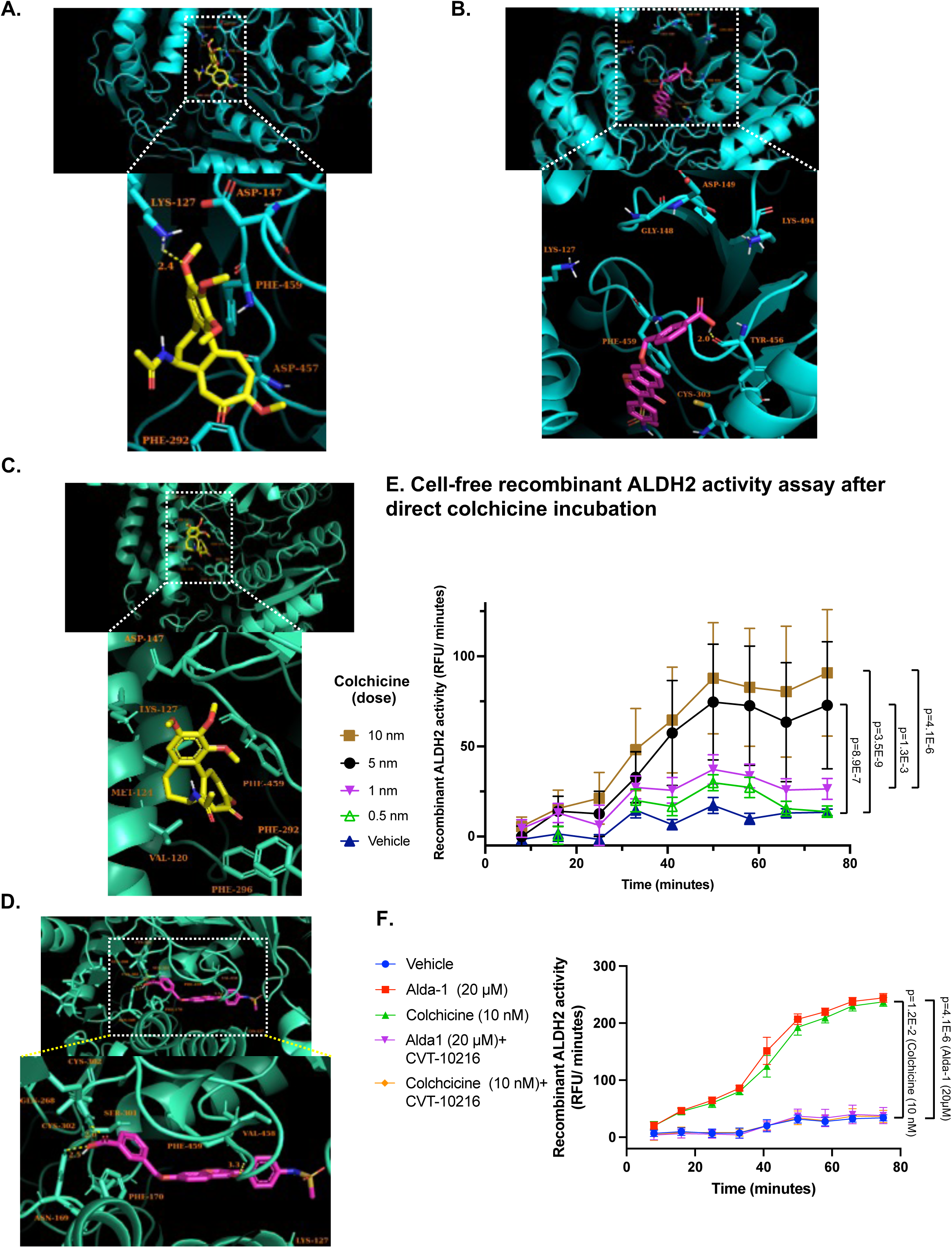
Predicted binding pose of colchicine and CVT-10216 within the 3INJ active site and 4FR8 site obtained by AutoDock Vina. (A) Predicted binding pose of colchicine within the 3INJ active site. Molecular docking of colchicine to the 3INJ was performed using AutoDock Vina v1.2.5 with the *vina* scoring function. The best-scoring pose (−6.427 kcal/mol) is shown as yellow sticks within the protein binding pocket rendered as cyan cartoon. Key interacting residues — Lys127, Asp147, Phe292, Asp457, and Phe459 — are displayed as sticks and labeled. A hydrogen bond is observed between the carbonyl oxygen of colchicine and Lys127 (2.4 Å, yellow dashed line). Additional stabilizing interactions are contributed by π–π stacking with Phe292 and hydrophobic contacts with Phe459. These interactions collectively stabilize colchicine in the active site cavity of 3INJ. Docking parameters: grid center = (6.365, −27.38, 44.879); grid size = 24 × 24 × 24 Å; grid spacing = 0.375 Å; exhaustiveness = 128. (B) Docking of CVT-10216 to the ALDH2 (3INJ) structure was performed using AutoDock Vina v1.2.5 with the *vina* scoring function. The best-scoring pose (−10.36 kcal/mol) is shown as magenta sticks within the cyan ribbon representation of the receptor. The ligand is positioned in the activator (vestibule) pocket, corresponding to the known Alda-1 binding site, and forms key interactions with residues Lys127, Asp147, Tyr249, Gly151, and Cys301. The lowest-energy conformation is stabilized by hydrogen bonds and hydrophobic interactions, with additional conformations (−10.31 to −10.15 kcal/mol) showing strong clustering (RMSD ≤ 2.5 Å), indicating good convergence. Docking parameters: grid center = (6.365, −27.38, 44.879); grid size = 24 × 24 × 24 Å; grid spacing = 0.375 Å; exhaustiveness = 128. (C) Docking of colchicine to the ALDH2 structure (PDB ID: 4FR8) was performed using AutoDock Vina v1.2.5 with the *vina* scoring function. The best-scoring pose (−6.665 kcal/mol) is displayed as yellow sticks, with the receptor shown in cyan ribbon representation. Colchicine is positioned in a hydrophobic pocket defined by Lys127, Asp147, Phe292, Val170, and Phe459, forming stabilizing hydrophobic and π–π interactions, particularly with Phe292 and Phe459. The lowest-energy conformation exhibited good clustering of poses (RMSD ≤ 3.5 Å), indicating consistent docking convergence. Docking parameters: grid center = (X = 1.43, Y = 49.958, Z = 49.385); grid size = 24 × 24 × 24 Å; grid spacing = 0.375 Å; exhaustiveness = 128. (D) Docking of CVT-10216 to ALDH2 (PDB ID: 4FR8). The best-scoring pose (−9.58 kcal/mol) is shown as magenta sticks within the cyan ribbon representation of the receptor. CVT occupies a hydrophobic cleft near the catalytic tunnel, forming hydrogen bonds with Cys302 (2.5 Å) and Ser301 (3.1 Å) and additional stabilizing interactions with Phe170, Val458, and Phe459. The ligand is further positioned in proximity to Glu268 and Lys127, residues known to contribute to the enzyme’s structural stability. The docking produced highly clustered top poses (RMSD ≤ 2 Å), indicating convergence on a stable binding mode. Docking parameters: grid center = (X = 1.43, Y = 49.958, Z = 49.385); grid size = 24 × 24 × 24 Å; grid spacing = 0.375 Å; exhaustiveness = 128. (E) Recombinant ALDH2 enzymatic activity was evaluated in a cell-free reaction system using the Abcam ALDH2 Activity Assay Kit, which quantifies NADH production in the presence of acetaldehyde and NAD⁺. Colchicine (0.5–10 nM) dose-dependently increased recombinant ALDH2 activity, demonstrating a direct activating effect of colchicine on the enzyme independent of cellular signaling or transcriptional regulation. (F) Comparison of recombinant ALDH2 enzymatic activity in a cell-free assay demonstrated that colchicine (10 nM) increased ALDH2 activity to a level comparable to that induced by the established ALDH2-specific activator Alda-1 (1 and 20 μM).

Similarly, CVT-10216 exhibited a strong binding preference for the same activator pocket. The best-ranked pose (−10.36 kcal/mol) was located within the Alda-1 binding region (**Fig. 6B**), engaging residues Lys127, Asp147, Tyr249, Gly151, and Cys301 through a combination of hydrogen bonding and hydrophobic interactions. The tight clustering of top poses (RMSD ≤ 2.5 Å) and the substantially lower predicted binding energy indicate a stronger interaction potential compared to colchicine. Together, these findings suggest that both colchicine and CVT-10216 can associate with the Alda-1 activator/vestibule pocket of ALDH2, though CVT-10216 demonstrates a higher predicted affinity and more extensive contact network. Therefore, CVT-10216 and colchicine could compete for binding to ALDH2 under certain conditions, with CVT-10216 likely having higher binding priority due to its stronger predicted affinity and more extensive contact profile. However, given potential differences in ligand size, flexibility, and solubility, the extent of in vivo competition would depend on their relative concentrations and access to the mitochondrial matrix where ALDH2 resides.

Docking simulations using AutoDock Vina v1.2.5 also revealed distinct yet adjacent binding sites for CVT-10216 and colchicine on the ALDH2 (4FR8) structure (**Fig. 6C, D**). CVT-10216 exhibited a strong predicted affinity (−9.58 kcal/mol) and bound deeply within the catalytic tunnel, forming hydrogen bonds with Cys302 (2.5 Å) and Ser301 (3.1 Å) and hydrophobic interactions with Phe170, Val458, and Phe459. These residues are located near the catalytic and cofactor-binding region, suggesting that CVT-10216 may interact competitively with the substrate or modulate catalytic accessibility (**Fig. 6D**).

In contrast, colchicine bound to a more peripheral hydrophobic pocket adjacent to the catalytic tunnel with a moderate predicted affinity (−6.67 kcal/mol) (**Fig. 6C**). The binding involved hydrophobic and π–π interactions with Phe292, Val170, and Phe459, and polar contacts with Lys127 and Asp147. This site partially overlaps with the Alda-1 activator pocket, consistent with experimental evidence that colchicine can activate ALDH2. The positioning of colchicine in this vestibule-like region suggests it may stabilize the enzyme in an active conformation through allosteric modulation rather than direct catalytic engagement.

Although CVT-10216 and colchicine share key hydrophobic contact residues (notably Phe170 and Phe459), docking to both 3INJ and 4FR8 indicates that the two ligands predominantly occupy the Alda-1 activator (vestibule) pocket of ALDH2 (**Fig. 6A-D**). Their binding orientations differ slightly: CVT-10216 extends more deeply toward the catalytic tunnel and forms additional contacts with Cys302 and Ser301, whereas colchicine remains closer to the outer region of the activator pocket, engaging Lys127, Asp147, and Phe292. This spatial overlap suggests that CVT-10216 and colchicine may compete for binding to the same allosteric pocket, with CVT-10216 likely exhibiting higher affinity and potential inhibitory effects, while colchicine may stabilize the enzyme in a more active conformation. Consequently, their opposing functional outcomes likely reflect differential engagement within a shared binding site rather than occupation of entirely distinct regions.

To determine whether colchicine directly activates ALDH2, we performed a cell-free in vitro enzymatic assay using purified recombinant ALDH2 protein (**Fig. 6E**). ALDH2 activity was quantified by measuring NADH production in the presence of acetaldehyde and NAD⁺ following direct incubation with colchicine. Colchicine increased recombinant ALDH2 activity in a concentration-dependent manner within the low nanomolar range, with higher concentrations producing greater NADH generation compared with vehicle control. Although the assay was not designed for formal kinetic analysis, the dose–response profile suggests an estimated half-maximal effective concentration (EC₅₀) of approximately 1–5 nM.

Notably, the reported EC₅₀ of the established ALDH2 activator Alda-1 is approximately 20–40 μM^28^, indicating that colchicine enhances ALDH2 activity at substantially lower concentrations. To further assess the relative potency of colchicine, we compared the effects of 20 μM Alda-1 and 10 nM colchicine on recombinant ALDH2 activity in a cell-free assay. Remarkably, 10 nM colchicine induced a level of ALDH2 activation comparable to that achieved with 20 μM Alda-1, suggesting that colchicine is a highly potent activator of ALDH2 in vitro.

To further examine whether colchicine and Alda-1 share a common interaction site on ALDH2, both compounds were co-incubated with the selective ALDH2 inhibitor CVT-10216. CVT-10216 completely abolished ALDH2 activation induced by either colchicine or Alda-1 in the cell-free assay, consistent with our in silico docking analysis predicting competitive binding of CVT-10216 with both compounds at ALDH2. Collectively, these findings demonstrate that colchicine directly enhances ALDH2 catalytic activity independent of cellular context and support a mechanism in which colchicine functions as a potent direct activator of ALDH2.

### Colchicine Attenuates IR-Induced Macrophage Senesence-Associated Stemness and Atherosclerosis in a Partial Left Carotid Ligation (PLCL) Model

To determine whether colchicine inhibits IR–induced atherosclerotic lesion formation, we employed our established partial left carotid ligation (PLCL) model^29^. This model was used to assess whether colchicine mitigates IR-mediated acceleration of atherosclerosis. As outlined in Fig. 7A, mice received AAV-PCSK9 and were placed on a high-fat diet (HFD) for 13 days prior to initiation of colchicine treatment (0.01 mg/kg/day, intraperitoneally). To minimize IR-associated body weight loss, mice were transitioned to a normal chow diet for two weeks following IR exposure and subsequently returned to HFD. One week after resuming HFD, PLCL surgery was performed, and carotid arteries were harvested two weeks later for analysis (**Fig. 7A**). Plasma cholesterol levels were significantly elevated in both colchicine- and vehicle-treated groups, with no significant difference between groups, indicating comparable hyperlipidemia (**Fig. 7B, and S1A**). Despite similar cholesterol levels, colchicine treatment markedly reduced atherosclerotic lesion size compared with vehicle controls both in male and female (**Fig. 7C, D and S1C**).

**Figure 7.**
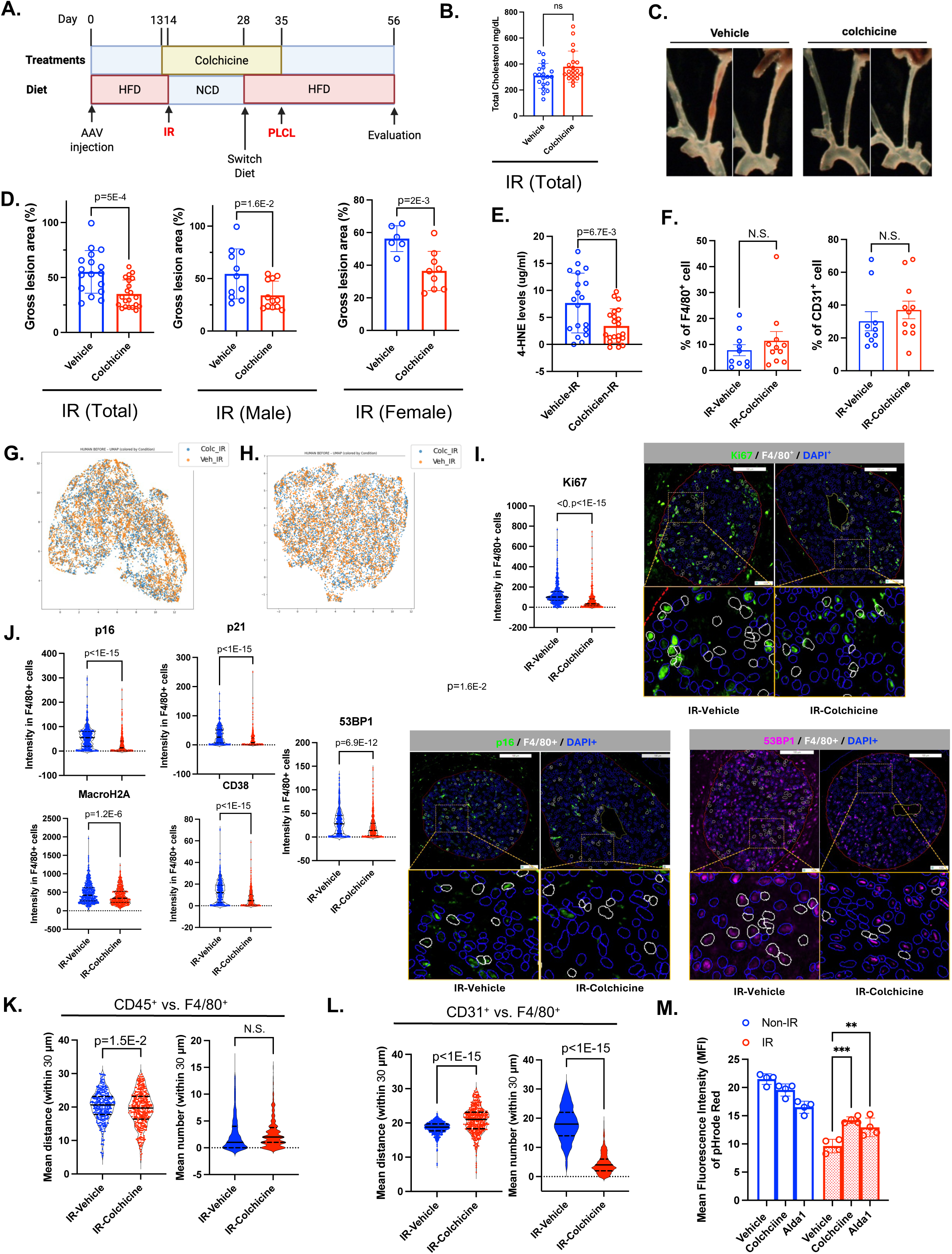
Colchicine inhibited IR-induced senescence and acceleration of atherosclerosis. **(A)** Experimental scheme of IR-induced acceleration of atherosclerosis and cholchicine treatment. **(B)** Cholesterol levels in mice treated with vehicle and colchicine. **(C, D)** Atherosclerosis lesions in vehicle and cholchicine group under IR. We evaluated the gross lesion area (%) as described in Methods (D). (E) Plasma 4-HNE level was measured as described in the methods. (F) Percentage of F4/80⁺ macrophages and CD31⁺ endothelial cells among total cells within the plaque for each group **(G–H)** UMAP visualization of single-cell data from vehicle- and colchicine-treated mice, demonstrating intermixing of cells and minimal batch effects. (**I, J**) Proliferation marker of Ki67 (**I**), and senescence-related molecules of p16, 53BP1, p21 (**S1D**), CD38 (**S1D**), and macroH2A (**S1D**) expression level (**J**) were was significantly lower in colchicine-treated mice compared to vehicle-treated mice.

To determine whether colchicine modulates ALDH2 loss–associated lipid aldehyde accumulation in vivo, plasma levels of the lipid peroxidation product 4-HNE were measured. Colchicine-treated mice exhibited significantly lower circulating 4-HNE levels compared with vehicle-treated controls (**Fig. 7E**). Because 4-HNE is a major toxic aldehyde detoxified by ALDH2, these data indicate a reduction in systemic aldehyde burden and are consistent with enhanced ALDH2-dependent detoxification in colchicine-treated mice.

To further characterize lesion composition, we performed COmbinatorial Microscopy Ensemble Technique (COMET) analysis with quantitative image analysis using the Visiopharm platform, enabling high-dimensional, spatially resolved single-cell protein profiling within tissue sections. Quantification of plaque composition showed no significant differences in the percentages of F4/80⁺ macrophages or CD31⁺ endothelial cells between colchicine- and vehicle-treated groups following IR (**Fig. 7F**). These results indicate that colchicine does not significantly affect inflammatory cell infiltration or plaque neovascularization after IR.

To assess potential batch effects in the single-cell RNA-sequencing analysis, Uniform Manifold Approximation and Projection (UMAP) embeddings were examined for vehicle-treated (n = 11; total cells = 5,011) and colchicine-treated mice (n = 12; total cells = 5,237). Cells from different experimental batches were well intermixed within each UMAP representation, with no evidence of batch-specific clustering (**Fig. 7G, H** and **S1B**), indicating minimal technical batch effects and preservation of the underlying biological structure. Given the robustness of these data and the substantial number of cells analyzed per group, downstream analyses were performed at the single-cell level rather than aggregated at the animal level, enabling high-resolution comparison of molecular expression across individual cells while minimizing the influence of inter-animal variability.

To determine the effects of colchicine on macrophage proliferation and stress responses, F4/80^+^ cells were analyzed for markers of proliferation, senescence, DNA damage, and apoptosis. COMET analysis demonstrated that colchicine significantly reduced expression of the proliferation marker Ki67, indicating suppression of macrophage proliferative activity (**Fig. 7I**). In parallel, colchicine markedly suppressed multiple markers of cellular senescence, including p21, p16, CD38, 53BP1, and MacroH2A, as quantified at the single-cell level (**Fig. 7J and S1D**). In addition, colchicine reduced expression of DNA damage response markers PARP1 and γH2AX, as well as the apoptosis marker caspase-7, indicating attenuation of DNA damage signaling and cell death pathways (**Fig. S2**). Collectively, these results demonstrate that colchicine limits macrophage proliferation while simultaneously suppressing cellular senescence, DNA damage responses, and apoptosis, consistent with an overall protective effect against IR-induced atherosclerosis formation.

Colchicine treatment also significantly reduced Ki67 expression and DNA damage markers in CD31^+^ endothelial cells and αSMA^+^ smooth muscle cells within plaques following IR exposure. However, unlike in macrophages, senescence-associated markers in CD31+ and αSMA+ cells were not uniformly decreased by colchicine treatment (**Fig. S4**). These findings suggest that the anti-senescent effects of colchicine are more broadly and consistently observed in macrophages, whereas in endothelial and smooth muscle cells, colchicine appears to modulate only selected senescence-associated programs rather than globally suppressing cellular senescence.

Colchicine decreased the spatial distance between CD45⁺ cells and F4/80⁺ macrophages, indicating that immune cell proximity within the plaque microenvironment was preserved (**Fig. 7K)**. In contrast, colchicine significantly increased the spatial distance between CD31⁺ cells and F4/80⁺ macrophages while reducing the number of CD31⁺–F4/80⁺ interactions, consistent with deeper localization of macrophages within the plaque. Although colchicine inhibited cell migration in our assay (**Fig. S2B**), efferocytosis was significantly enhanced following treatment (**Fig. 7M**). In parallel, colchicine increased intracellular NAD⁺ and ATP levels (**Fig. 2C,D**), which may enhance macrophage efferocytic capacity, facilitate deeper macrophage localization within atherosclerotic plaques, and reduce the accumulation of DNA-damaged (PARP1^hi^ and γH2AX^hi^) and apoptotic (caspase-7^hi^) cells (**Fig. S3A**). Additionally, colchicine decreased macrophage expression of the anti-phagocytic “don’t eat me” signal SIRPα, further suggesting enhanced clearance of senescent macrophages through efferocytosis (**Fig. S3A**). This mechanism may contribute to the reduced burden of apoptotic macrophages within the plaque and the attenuation of vulnerable plaque formation.

Furthermore, spatial single-cell analysis demonstrated strong linear correlations between the senescence markers p16 and p21 and Ki67 in the majority of F4/80⁺ macrophages within plaques following IR exposure (**Fig. 8A,B,D**). As we previously reported in murine plaque spatial single-cell analyses, a subset of cells exhibiting a linear relationship between senescence markers and Ki67 represents a senescence-associated stemness (SAS) state^30^. In the present study, IR exposure markedly strengthened this linear correlation (**Fig. 8B, D**), indicating an expansion of the SAS-like macrophage population within plaques. This phenotype is characterized by persistent expression of senescence markers despite retained proliferative potential, suggesting escape from senescence-associated cell-cycle arrest. Notably, colchicine treatment shifted the p16/p21–Ki67 correlation toward the lower end of the slope (**Fig. 8C, E**), consistent with suppression of the SAS-like phenotype in plaque macrophages.

**Figure 8.**
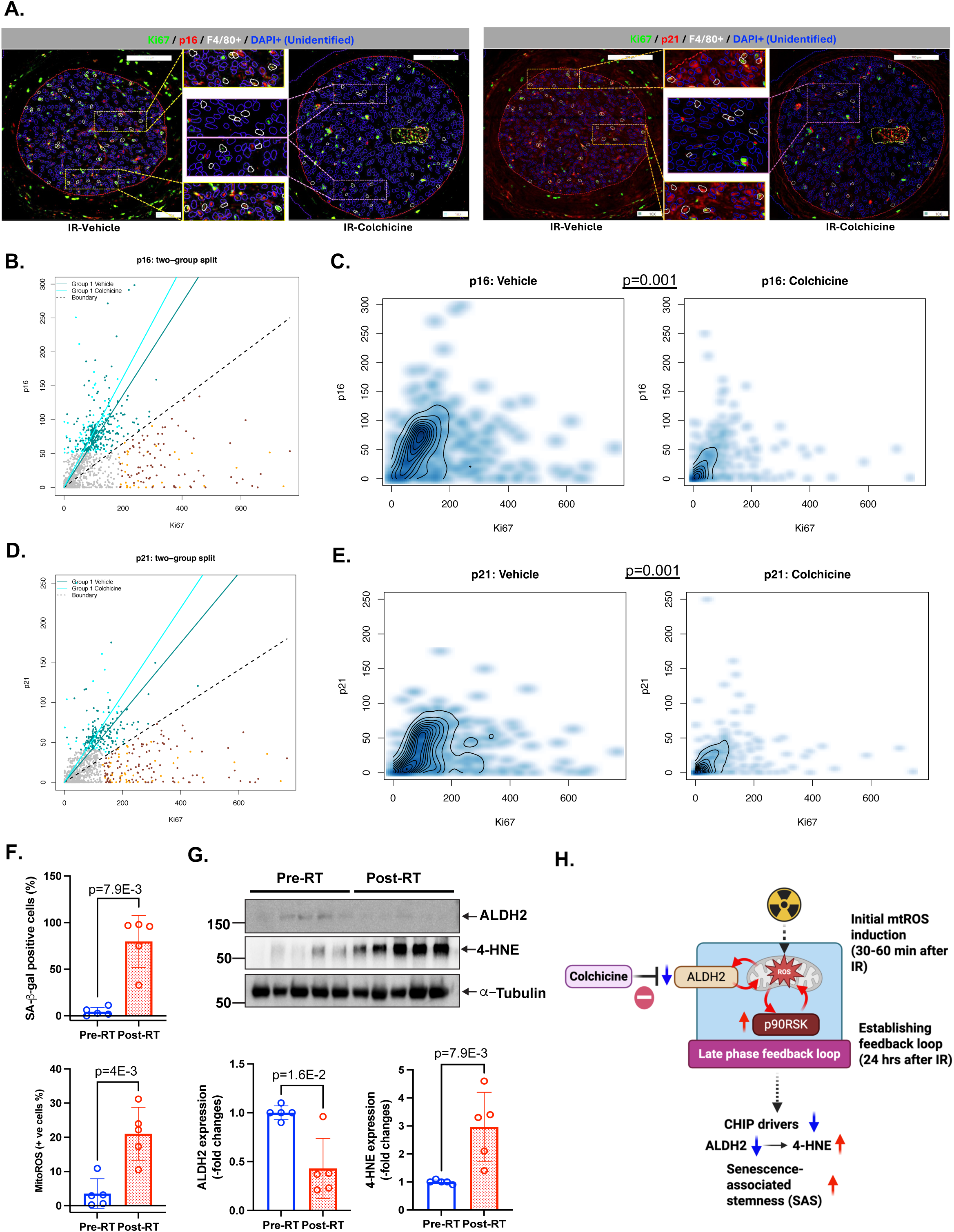
Colchicine Inhibited IR-Induced Senescence-Associated Stemness (SAS). **(A-E)** Spatial single-cell analysis revealed strong linear correlations between the senescence markers p16, and p21 and the proliferation marker Ki67 in F4/80⁺ cells within plaques following IR exposure (**B, D**). Representative images corresponding to the quantified markers are shown in (**A**). (**C, E**) Two-dimensional density plots of Ki67 versus p16 (**C**) and Ki67 versus p21 (**E**) expression in F4/80⁺ macrophages from vehicle- and colchicine-treated groups. Colchicine treatment reduced the accumulation of p16⁺ and p21⁺ cells and decreased their overlap with proliferative (Ki67⁺) populations. Differences in the overall joint distributions between groups were evaluated using a nonparametric two-sample energy distance test with 999 permutations; corresponding p values indicate the statistical significance of distributional differences between vehicle and colchicine-treated cells. **(F, G)** Human monocyte-derived macrophages (HMDMs) generated from PBMCs of five patients (Table S2) collected before radiation therapy and 3–6 months after radiation therapy were analyzed for SA-β-gal activity (F, top), mitochondrial ROS production (F, bottom), and protein expression by Western blotting using the indicated antibodies (G, top) and quantification (G, bottom). **(H) Graphical abstract:** IR inhibits ALDH2, although ALDH2 does not contribute to the early phase of the response. Initially, IR induces mtROS within 30-60 minutes, which activates p90RSK and triggers a positive feedback loop, as previously reported^18^. This feedback loop contributes to the subsequent activation of a late-phase process. The inhibition of ALDH2 by IR plays a key role in this process, driving the reduction of CH drivers, the induction of senescence, and the accumulation of 4-HNE approximately 24 hours after IR exposure. These changes promote cellular aging and the development of atherosclerosis. By targeting ALDH2, colchicine disrupts this feedback loop, preventing the reduction of CH drivers and inhibiting the progression of senescence, including its functional unique type of senescence-associated stemness (SAS).

Collectively, these findings indicate that colchicine mitigates IR-induced atherosclerosis through coordinated suppression of systemic lipid peroxidation, macrophage senescence, DNA damage signaling, immune cell infiltration, efferocytic activity, and the SAS-like phenotype within plaques, ultimately reducing plaque burden and favoring plaque stabilization.

### Radiation Therapy Induces 4-HNE Accumulation and Senescence in Human Monocyte-Derived Macrophages (HMDMs) obtained from esophagus and lung cancer patients treated by radiation

To determine the effects of radiation therapy on the ALDH2–4-hydroxynonenal (4-HNE) axis in humans, peripheral blood mononuclear cells (PBMCs) were collected from patients with lung or esophageal cancer before and after thoracic radiation therapy (**Table S2**) and differentiated into human monocyte–derived macrophages (HMDMs). ALDH2 expression, 4-HNE accumulation, mitochondrial reactive oxygen species (mtROS) production, and senescence-associated β-galactosidase (SA–β-gal) activity were then assessed. Compared with pre-radiation samples, HMDMs obtained after radiation therapy exhibited a significant increase in 4-HNE accumulation and mtROS production, accompanied by a marked increase in SA–β-gal–positive cells. In contrast, ALDH2 expression was significantly reduced in post-radiation HMDMs. These changes were consistently observed across patient samples. Collectively, these results demonstrate that thoracic radiation therapy is associated with suppression of ALDH2 and activation of oxidative stress and senescence pathways in human macrophages, indicating that ALDH2 dysregulation occurs not only in metabolic disease states^31^ but also following radiation exposure in humans.

## Discussion

In the present study, we demonstrate that exposure to IR induces a SAS program in plaque macrophages, characterized by acquisition of stemness-related features within the atherosclerotic plaque microenvironment. This IR-induced SAS occurred in parallel with senescence-associated inflammatory signaling and reflects a non-canonical senescence state in which cells exhibit durable stress responses while adopting stemness-associated gene programs, rather than simple proliferative arrest. The emergence of SAS within plaques after IR exposure provides a potential mechanistic link between IR exposure and persistent plaque inflammation and remodeling. Colchicine markedly suppressed IR-induced SAS in plaque macrophages. We identify direct activation of ALDH2 by colchicine as a previously unrecognized and central mechanism underlying its protective effects against IR-accelerated atherosclerosis. Through ALDH2 activation, colchicine suppresses mitochondrial oxidative stress, restores CH driver expression, and limits acquisition of stemness-associated transcriptional programs. Consistent with this mechanism, RNA sequencing of myeloid cells revealed that colchicine broadly reprograms IR-induced transcriptional responses, prominently modulating NRF2 and senescence-related gene signatures (**Fig. 1, and 2**).

Building on our prior work showing that IR-induced mtROS activate p90RSK to prime macrophages for exaggerated secondary oxidative stress responses^18^, the current study further uncovers a positive feedback loop between mtROS and p90RSK activation in IR-exposed macrophages. Within this loop, ALDH2 emerges as a critical modulator of sustained oxidative stress and senescence signaling. Direct activation of ALDH2 by colchicine reduced mtROS accumulation, limited lipid peroxidation as reflected by decreased 4-HNE levels, and reversed IR-mediated suppression of key CH drivers, including TET2 and DNMT3A (**Fig. 3, and 4**). These effects were accompanied by robust inhibition of SAS induction in plaque macrophages.

Mechanistically, pharmacological inhibition of mtROS with mitoTEMPOL or p90RSK with FMK-MEA significantly attenuated IR-induced downregulation of ALDH2 and CH drivers, identifying mtROS–p90RSK signaling as a key contributor to senescence development following IR exposure (**Fig. 3**). Notably, colchicine did not alter early-phase IR-induced mtROS generation or initial p90RSK activation (**Fig. 3H-L**), indicating that its protective effects occur downstream of the initial oxidative insult. Together, these findings support a model in which IR-induced mtROS–p90RSK signaling promotes senescence within atherosclerotic plaques through suppression of ALDH2, and in which colchicine interrupts this pathogenic program by directly restoring ALDH2 activity.

The docking analyses and structural visualizations support a model in which colchicine functions as an allosteric modulator of ALDH2 rather than a catalytic inhibitor. Notably, colchicine’s predicted binding site overlaps with that of CVT-10216, a well-characterized ALDH2 inhibitor, suggesting that the two compounds may compete for occupancy of the same regulatory pocket (**Fig. 6**). Whereas CVT-10216 destabilizes ALDH2 and suppresses its enzymatic activity, colchicine binding appears to enhance ALDH2 function. This divergence in functional outcomes despite competition for a shared binding site highlights a previously underappreciated pharmacological interaction. Consistent with this model, we demonstrate for the first time that colchicine directly activates ALDH2 enzymatic activity in vitro. Pretreatment with colchicine rendered CVT-10216 ineffective (**Fig. 5F-H**), whereas pretreatment with CVT-10216 significantly attenuated colchicine-mediated ALDH2 activation and inhibited IR-induced 4-HNE accumulation (**Fig. 5A–C**). This reciprocal interference strongly suggests competitive binding or regulation at overlapping ALDH2 sites, a conclusion further supported by molecular docking and structural modeling. Collectively, these findings identify ALDH2 as a novel direct molecular target of colchicine. Through ALDH2 activation, colchicine disrupts the IR-induced ALDH2–4-HNE–mtROS–p90RSK positive feedback loop, thereby attenuating radiation-induced cellular senescence (**Fig. 8H**). Importantly, this mechanism is independent of colchicine’s established effects on microtubule dynamics and reveals a distinct pathway by which colchicine mitigates persistent oxidative stress and premature senescence following cancer therapy.

Importantly, ALDH2 expression is frequently downregulated in various tumor types and is often associated with clinical outcomes and immune signatures. In certain cancers, such as gastric cancer, higher ALDH2 expression correlates with improved survival, consistent with its role in aldehyde detoxification and tumor suppression^32,33^. These expression and epigenetic patterns are observed across diverse ancestries and are not solely attributable to the rs671 polymorphism^33^. Furthermore, in a nationwide propensity-matched cohort of patients with immune-mediated inflammatory diseases, colchicine use was linked to decreased all-cancer and colorectal cancer incidence^34^. Taken together, our data suggest that ALDH2 activation by colchicine offers dual therapeutic benefits: it may protect against cardiovascular complications following cancer therapy and also improve cancer prognosis by suppressing SAS. This beneficial effect is independent of colchicine’s anti-inflammatory action via microtubule disruption, and instead stems from its ability to modulate mitochondrial oxidative stress and senescence pathways through ALDH2 activation.

We found that IR inhibited DNMT3a and TET2 expression. It is well known that CH mutation can change CH drivers function and increase ROS production, which subsequently induce vascular inflammation and atherosclerosis. In this study, we found that low dose of 2-5 Gy of radiation inhibited DNMT3a and TET2 expression by promoting protein degradation in myeloid cells. Since CH mutation of DNMT3a and TET2 inhibit those two-enzyme function and accelerate atherosclerosis formation have been reported^35^, the reduction of DNMT3a and TET2 expression after radiation in macrophage may play a role in RICVD. In this study, we found the contribution of mtROS production and p90RSK activation in regulating radiation-induced CH drivers protein levels reduction, but the exact molecular mechanism remains unclear, and further investigation will be necessary.

In summary, we identify direct activation of ALDH2 as a central and previously unrecognized mechanism underlying colchicine’s protective effects against IR-induced vascular injury. By restoring ALDH2 activity, colchicine suppresses mtROS–p90RSK signaling, preserves cholesterol homeostasis, and limits stress-associated senescence, which is particularly enhanced following IR exposure, thereby attenuating atherosclerosis progression. These findings establish ALDH2 as a novel and essential molecular target of colchicine and provide a strong mechanistic foundation for therapeutic strategies aimed at ALDH2 in radiation-induced vascular disease and aging-related pathologies. Importantly, this mechanism extends beyond colchicine’s known anti-inflammatory actions and may offer a new approach to reducing cardiovascular disease risk in cancer survivors without compromising immune surveillance. Moreover, given the prognostic relevance of ALDH2 in multiple cancers, its activation may also contribute to improved cancer outcomes.

## Funding Sources

This work was partially supported by grants from the National Institutes of Health (NIH) to Dr. Abe (HL-149303, HL-179815, and AI-156921), Dr. Cooke (HL-148338, HL-133254, HL-157790, and HL-149303), and Dr. Le (HL-134740 and HL-149303). This study was also partially supported by the University of Texas MD Anderson Cancer Center Institutional Research Grant (IRG) Program awarded to Dr. Kotla. Research was performed in the Flow Cytometry and Cellular Imaging Core Facility at The University of Texas MD Anderson Cancer Center, which is supported in part by the National Institutes of Health through MD Anderson’s Cancer Center Support Grant CA016672, the National Cancer Institute Research Specialist Award R50 CA243707-01A1, and a Shared Instrumentation Award from the Cancer Prevention Research Institute of Texas (CPRIT; RP121010).

## Contributions

J.A. planned and generated the study design, obtained funding, interpreted data, and wrote the manuscript. V.K.S. and J.L. performed the experiments, interpreted the data, and wrote the manuscript. W.C. supported data analyses, and wrote the manuscript. G.F.M., K.A.K., A.K.R., performed the surgery and the following analysis. O.H. performed the experiments and maintained mouse colonies. L.A.R., K.Y.C., J.H.K., S.F.L.M., S.A., K.O.M., E.G.S. M.O. performed the experiments and analyzed the data. S.L. supported data analyses. A.D., J.H., J.P.C., K.L.S., K.F., L. Y-C., C.M., P.A., S.W.Y., R.P., J.K.B., N.L.P., K.T.N., M.H., C.D.F., E.K., and S.K. contributed to the interpretation of the data. S.H.L., G.W., N-T. L., and S.K. planned and generated the study design, performed clinical study, obtained funding, interpreted data, and wrote the manuscript.

## Disclosures

S.H.L is an Advisory Board member of AstraZeneca, Beyond Spring Pharmaceuticals, STCube Pharmaceuticals. The other authors report no conflicts.

## Supplemental Materials

Supplementary Methods

Contact for Reagent and Resource Sharing

Experimental model and Subject Details

Methods Details

Online Supplementary Figures S1-S4

Online Supplementary Figure Legends S1-4

Online Supplementary Table S1-2

Raw Data Sets File1: Intensity of COMET analysis

Raw Data Sets File 2: Distance between CD31^+^ and F4/80^+^ cells

Raw Data Sets File 3: Distance between CD4^+^/CD8^+^/CD45^+^ and F4/80^+^ cells

Uncropped Western Blots

## Non-standard Abbreviations and Acronyms

ALDH2: aldehyde dehydrogenase 2
AAV: adeno-associated virus
ANOVA: analysis of variance
ASCVD: atherosclerotic cardiovascular disease
ATF4: activating transcription factor 4
BMDMs: bone marrow–derived macrophages
BMI: body mass index
CH: clonal hematopoiesis
CHIP: clonal hematopoiesis of indeterminate potential
COMET: COmbinatorial Microscopy Ensemble Technique
CRP: C-reactive protein
CVD: cardiovascular disease
DDR: DNA damage response
GO: gene ontology
GSEA: gene set enrichment analysis
HFD: high-fat diet
HMDMs: human monocyte–derived macrophages
HIPAA: Health Insurance Portability and Accountability Act
IACUC: Institutional Animal Care and Use Committee
IMC: imaging mass cytometry
IR: ionizing radiation
LDL: low-density lipoprotein
MACE: major adverse cardiovascular events
mtROS: mitochondrial reactive oxygen species
NRF2: nuclear factor erythroid 2–related factor 2
PBMCs: peripheral blood mononuclear cells
PLCL: partial left carotid ligation
qRT-PCR: quantitative reverse-transcription polymerase chain reaction
RICVD: radiation-induced cardiovascular disease
ROI: region of interest
RT: radiation therapy
SA-β-gal: senescence-associated β-galactosidase
SAS: senescence-associated stemness
SASP: senescence-associated secretory phenotype
UMAP: Uniform Manifold Approximation and Projection
4-HNE: 4-hydroxynonenal

